# Design of Peptide-Based Protein Degraders via Contrastive Deep Learning

**DOI:** 10.1101/2022.05.23.493169

**Authors:** Kalyan Palepu, Manvitha Ponnapati, Suhaas Bhat, Emma Tysinger, Teodora Stan, Garyk Brixi, Sabrina R.T. Koseki, Pranam Chatterjee

**Affiliations:** Department of Biomedical Engineering, Duke University; Harvard University; Center for Bits and Atoms, MIT; MIT Media Lab

## Abstract

Therapeutic modalities targeting pathogenic proteins are the gold standard of treatment for multiple disease indications. Unfortunately, a significant portion of these proteins are considered “undruggable” by standard small molecule-based approaches, largely due to their disordered nature and instability. Designing functional peptides to undruggable targets, either as standalone binders or fusions to effector domains, thus presents a unique opportunity for therapeutic intervention. In this work, we adapt recent models for contrastive language-image pre-training (CLIP) to devise a unified, sequence-based framework to design target-specific peptides. Furthermore, by leveraging known experimental binding proteins as scaffolds, we create a streamlined inference pipeline, termed **Cut&CLIP**, that efficiently selects peptides for downstream screening. Finally, we experimentally fuse candidate peptides to E3 ubiquitin ligase domains and demonstrate robust intracellular degradation of pathogenic protein targets in human cells, motivating further development of our technology for future clinical translation.

## Introduction

It has been estimated that while nearly 15% of human proteins are disease-associated, only 10% of such proteins interact with currently-approved small molecule drugs. Even more strikingly, of the 650,000 protein-protein interactions (PPIs) in the proteome, only 2% are considered “druggable” by pharmocological means [Shin et al., 2020]. Aside from small molecule-based approaches, monoclonal antibodies have found significant success in the clinic as biologics, but while they are highly selective and can bind antigens with high specificity, they are limited to extracellular PPIs and cannot naturally permeate the cell membrane [Slastnikova et al., 2018].

Peptides have been widely recognized as a more selective, effective, and safe method for targeting pathogenic proteins, due to their sequence-specific binding to regions of partner molecules [Padhi et al., 2014, Buchwald et al., 2014]. They have further demonstrated targeting of both extracellular and intracellular proteins, due to their small size and enhanced permeability, with or without conjugation to cell penetrating peptide (CPP) sequences [Lindgren et al., 2000, Lozano et al., 2017, Adhikari et al., 2018]. Beyond standalone peptide binders, our group has recently fused computationally-designed peptides to effector domains, such as E3 ubiquitin ligases, to enable binding and selective intracellular degradation of pathogenic targets of interest [Chatterjee et al., 2020]. Extending this “ubiquibody” (uAb) strategy to undruggable targets, including numerous oncogenic and viral proteins, represents a promising new therapeutic approach.

Current approaches for peptide engineering have relied on high-throughput screening and structure-based rational design, with the goal of redirecting to alternate targets, extending half-life *in vivo*, improving solubility, or preventing aggregation [Fosgerau and Hoffmann, 2015]. Experimental methods, such as large phage display libraries and quantitative binding assays, while effective at selecting strong candidate sequences, are laborious and expensive [Wu et al., 2016, Kong et al., 2020, Carle et al., 2021]. Structure-based methods for peptide design consist of interface predictors and peptide-protein docking softwares [Raveh et al., 2011, Sedan et al., 2016, Tsaban et al., 2022]. These approaches, however, rely heavily on the existence of co-crystal complexes consisting of the target protein, thus excluding disordered or unstable proteins, such as transcription factors, which have significant disease implications and are difficult to solve via experimental or computational protein structure determination methods [Peterson et al., 2017, Das et al., 2018, Jumper et al., 2021]. Therefore, there is a need for the development of a sequence-based peptide generation platform, so as to rapidly and programmbly design peptides to any target protein, especially those for which minimal structural information exists.

The sequential structure of proteins, along with their hierarchical semantics, makes them a natural target for language modeling. Recently, language models have been pre-trained on over 200 million natural protein sequences to generate latent embeddings that grasp relevant physicochemical, functional, and most notably, tertiary structural information [Rives et al., 2021, Elnaggar et al., 2020, Vig et al., 2020, Rao et al., 2020]. Even more interestingly, generative protein transformers have produced novel protein sequences with validated functional capability [Madani et al., 2021]. Additionally, by augmenting input sequences with their evolutionarily-related counterparts, in the form of multiple sequence alignments (MSAs), the predictive power of protein language models can be further strengthened, as demonstrated by state-of-the-art contact prediction results [Rao et al., 2021].

In this study, we combine pre-trained protein language embeddings with novel contrastive learning architectures for the specific task of designing peptide sequences that bind target proteins and induce their degradation when fused to E3 ubiquitin ligase domains. By jointly training protein and peptide encoders to capture similarities between known peptide-protein pairs, our model accurately evaluates peptide inputs as potential binders for embedded target proteins. To further downselect initial peptide candidate lists for queried targets, we use predicted or experimentally-validated binding proteins as scaffolds for splicing, thus creating an integrated inference pipeline known as Cut&CLIP. We demonstrate that Cut&CLIP reliably and efficiently generates peptides which, when experimentally integrated within a uAb construct, induce robust degradation of pathogenic proteins in human cells.

## Results

### Dataset Curation and Augmentation

Previously, we demonstrated the utility of using scaffold proteins to derive functional peptides for uAb generation by executing the PeptiDerive protocol on co-crystals containing the target protein, thus identifying the linear polypeptide segments suggested to contribute most to binding energy [Chatterjee et al., 2020, Sedan et al., 2016]. To create a comprehensive dataset of these computationally-derived presumptive peptides, we applied PeptiDerive to complexes in the Database of Interacting Protein Structures (DIPS) [Sedan et al., 2016, Townshend et al., 2018]. We ran PeptiDerive on every co-crystal in DIPS with a resolution of ≤ 2 Å, and selected the top 20-mer peptides for each to include in the dataset, generating a total of 28,517 peptide-receptor pairs. Finally, we appended the Propedia dataset, an experimentally-derived database that includes an additional 19,814 peptide-receptor complexes from the Protein Data Bank (PDB) [Martins et al., 2021]. Together, our database represents one of the most comprehensive collections of peptide-protein pairs, and can thus serve as a standardized training set for future interface modeling.

For training, we specifically clustered the protein receptor sequences at 50% sequence identity using MMSeq2, yielding 7434 clusters, and split the clusters into train, validation, and test splits according at a 0.7/0.15/0.15 ratio, respectively [Steinegger and Söding, 2018]. All sequences from the selected clusters were used in the train and validation splits, but only a single representative sequence for each cluster was employed for the test split in order to ensure a reasonable balance of sequence diversity.

### Model Architecture and Training

One of the core problems in computer vision and natural language processing (NLP) is zero-shot classification. In the last year, OpenAI has introduced a new architecture, termed CLIP (Contrastive Language-Image Pre-Training), which utilizes zero-shot transfer and multimodal learning to associate visual concepts in images and link them with their captions [Radford et al., 2021]. Thus, we hypothesized that just as CLIP connects images to their corresponding captions using jointly-trained image and caption encoders, we can leverage CLIP-based architecture to map target proteins to their corresponding peptides using jointly trained receptor and peptide encoders. To do this, we trained encoders such that the cosine similarity between receptor embeddings and peptide embeddings, defined as

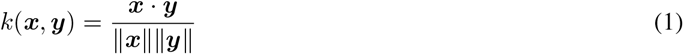

is near 1 for receptor-peptide pairs which do bind to each other, and is near -1 for receptor-peptide pairs which do not bind to each other. As input, the receptor encoder uses an MSA, while the peptide encoder simply uses the peptide sequence.

Next, the receptor and peptide encoders were trained on batches of *n* pairs of receptors and peptides which are known to interact. Receptor MSAs and peptide sequences were encoded by their respective encoders, producing receptor embeddings ***r***_**1**_, …, ***r***_***n***_, and peptide embeddings ***p***_**1**_, …, ***p***_***n***_. The cosine similarity between all *n*^2^ receptor and peptide pairs is computed in a matrix ***K***, defined such that

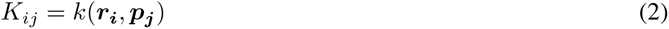

We then interpreted these cosine similarities as softmax logits. Logits were scaled by a learned temperature parameter *t*, which controls the model’s degree of uncertainty in output probabilities [Hinton et al., 2015]. We define two cross-entropy losses, one on the matrix rows and one the matrix columns:

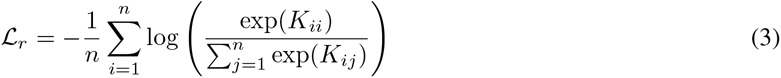

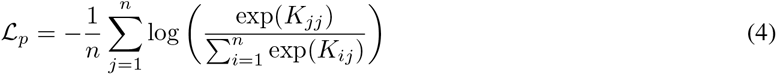

*ℒ*_*r*_ represents the loss on the model’s ability to predict the correct receptor given a single peptide, while *ℒ*_*p*_ represents the loss on the model’s ability to predict the correct peptide given a single receptor. By using these cross-entropy losses, we implicitly assumed that the *n*^2^ − *n* receptor-peptide pairs in the batch which are not known interactions do not bind at all. While this may not be a completely accurate assumption, it is approximately true. The model was then trained on the average of these two losses. The entire training process is illustrated in Figure 1.

**Figure 1:**
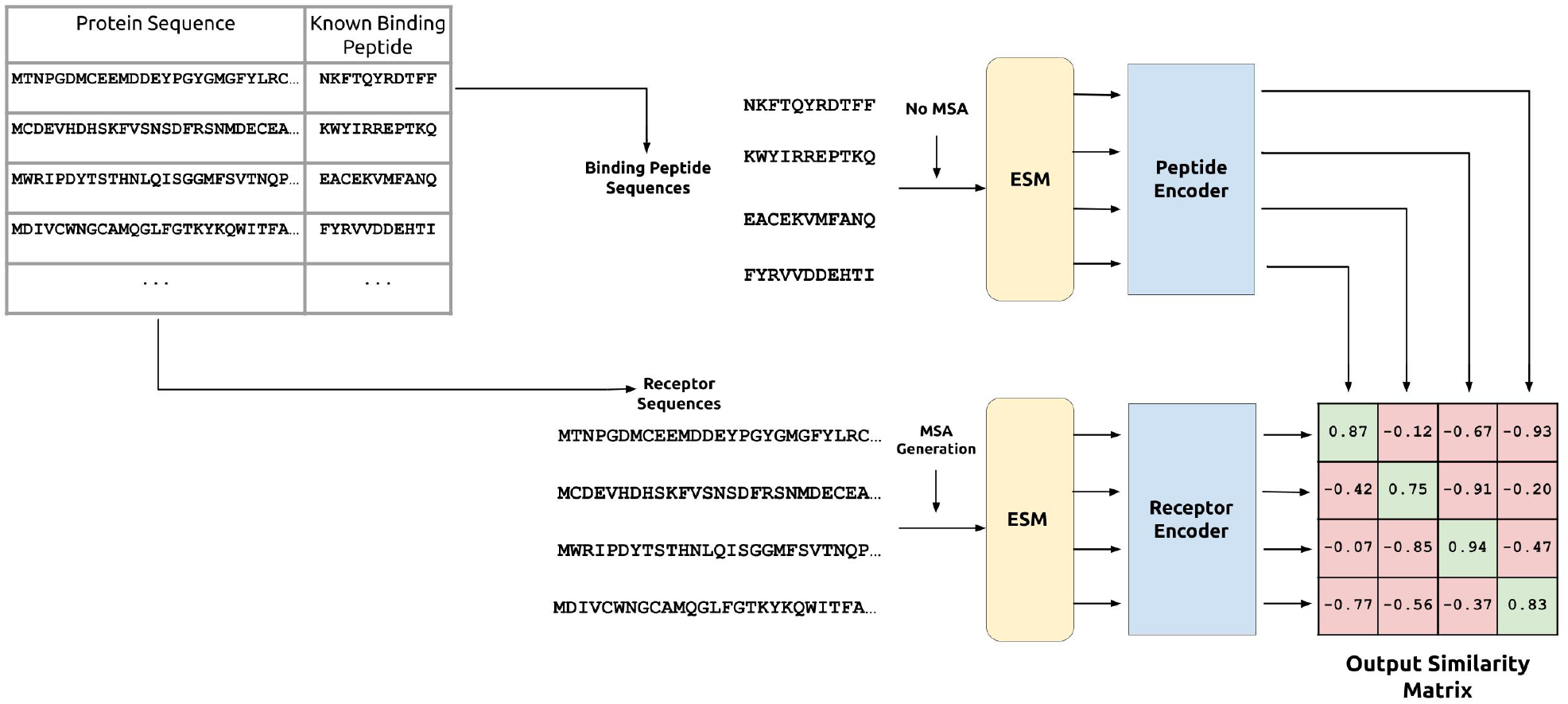
CLIP training process for peptide-protein pairs. Peptide and receptor encoders are jointly trained on ESM embeddings to predict high cosine similarities between known peptide-receptor embedding pairs and low cosine similarities for all other pairs.

Specifically, receptor MSAs and peptide sequences were first input into the ESM pre-trained transformer protein language models introduced previously by Facebook [Rives et al., 2021, Rao et al., 2020]. These pre-trained models were trained on millions of diverse amino acid sequences, allowing our encoders to extract feature-rich embeddings, which are robust to sequence diversity while being trained on a relatively small dataset. We employed the ESM-MSA-1b model for the receptor MSAs, and ESM-1b for the peptide sequences, which does not require MSA inputs (Figure 1). We then trained the receptor and peptide encoders taking these ESM embeddings as input.

The receptor encoder and peptide encoder have identical architectures, though they differ in hyperparameters such as the number of layers. Starting with an input *l* × *e*_*i*_ ESM embedding (where *l* is the input sequence length and *e*_*i*_ is the dimension of the ESM embedding), we applied *h*_1_ feedforward layers with ReLU activation on each amino acid embedding separately, producing a *l* × *e*_*o*_ embedding, where *e*_*o*_ is the output embedding dimension produced by our encoder. Then, we then averaged the embedding over the length dimension, producing an embedding vector of length *e*_*o*_. Finally, we applied *h*_2_ feedforward layers with ReLU activation on the embedding vector to get the output embedding. Notably, since the first set of hidden layers operate on single amino acids in isolation, and the second set of hidden layers operate on the embedding averaged over the length dimension, the encoder has minimal dependence on the sequence length. This is particularly important for helping the peptide encoder generalize to a broad range of peptide lengths.

As a relevant metric for model assessment in the context of screening, we calculated the top-*k* accuracy, which is the probability that the correct peptide is in the top *k* when provided a fixed batch of 250 candidate peptides, a suitable threshold for genetic screening. To calculate this metric, during prediction, we provide the model with a single protein target receptor and 250 peptides from our training set, one of which is a known binder.

Post-training, our final models demonstrate accurate ranking of known targeting peptides for a given target and vice versa, achieving 50% probability of identifying a correct candidate in the ranked top 50 out of 250, for example. These results motivate not only model application for tractable peptide screening assays, but also its utilization to evaluate peptide specificity to a desired target, in comparison to off-target receptor proteins (Figure 2).

**Figure 2:**
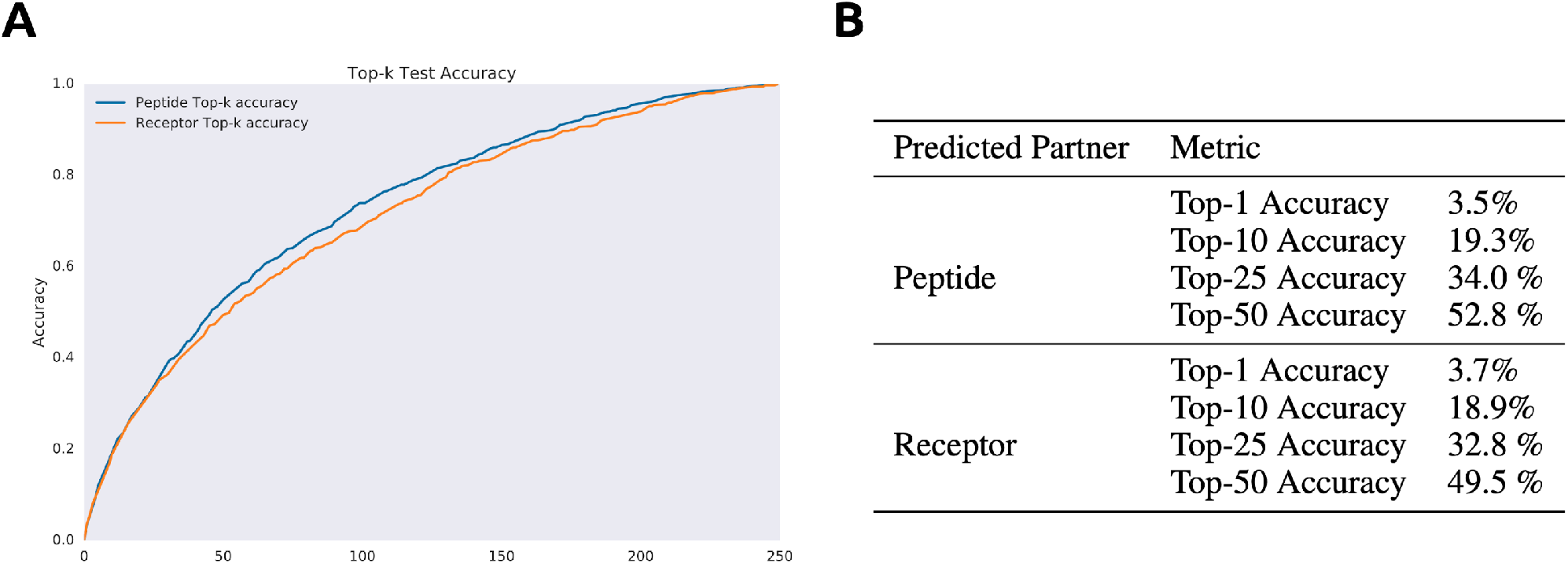
Results of model validation and testing. A) Top-*k* accuracy of predicting the correct binding partner out of a batch of 250. B) Selected test results. Accuracies are calculated via selection of the known binding partner out of a batch of 250 to a queried target. All results from an ensemble of the top four trained models.

### Cut&CLIP Inference

Inspired by our previous results leveraging co-crystals of target proteins and known binders to design effective peptide-based degraders, we decided to employ our CLIP model to predict binding peptides using experimentally-validated interacting proteins for a queried target [Chatterjee et al., 2020]. As opposed to our previous work, the current inference pipeline simply requires the sequence of potential binders from established PPI databases or experimental screening results, enabling us to be more flexible in identifying starting scaffolds [Szklarczyk et al., 2020, Johnson et al., 2021]. We then compute the CLIP peptide embedding for all *k*-mers of the interacting protein (where *k* is the desired size of our peptide), and rank them by their cosine similarities with the CLIP receptor embedding of the target protein. Our final Cut&CLIP pipeline is illustrated in Figure 3.

**Figure 3:**
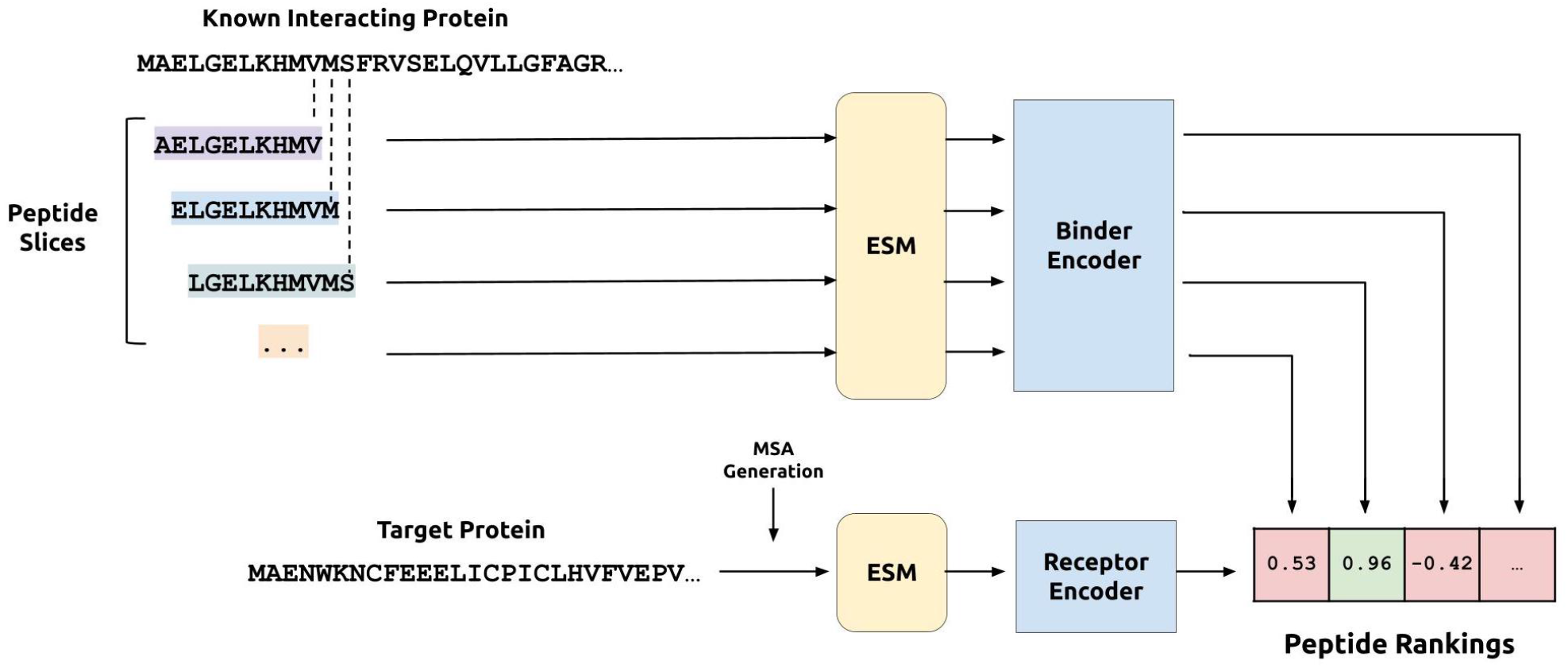
The Cut&CLIP inference protocol. A known interacting protein which is validated to interact with the target protein is cut up into peptide-size slices, enabling downstream ranking via the trained CLIP model.

While this strategy may not be sufficient to identify stand-alone peptide binders with high target affinity, our motivation derives from the well-understood catalytic nature of E3 ubiquitin ligase activity, where selective target binding is sufficient to induce degradation [Békés et al., 2022, Buetow and Huang, 2016, Portnoff et al., 2014].

### Target Protein Degradation with Cut&CLIP-Derived Peptides

Numerous previous works have attempted to redirect E3 ubiquitin ligases by replacing their natural protein binding domains with those targeting specific proteins [Gosink and Vierstra, 1995, Zhou et al., 2000, Su et al., 2003]. Recently, based on the seminal work of Portnoff, et al., our group reprogrammed the specificity of a modular human E3 ubiquitin ligase called CHIP (carboxyl-terminus of Hsc70-interacting protein) by replacing its natural substrate-binding domain, TPR, with designer peptides to generate minimal and programmable uAb architectures [Portnoff et al., 2014, Chatterjee et al., 2020]. To evaluate Cut&CLIP’s utility as compared to a slower structure-based method using AlphaFold [Jumper et al., 2021], we selected three target proteins for experimental characterization: the spike receptor binding domain (RBD) of SARS-CoV-2, the TRIM8 E3 ubiquitin ligase, and the KRAS oncoprotein.

Previously, we demonstrated robust degradation of RBD using peptide-based uAbs, and with stable co-crystal structures of RBD and the human ACE2 receptor, it represents a very tractable target for standard structure-based peptide generation [Chatterjee et al., 2020, Lan et al., 2020]. TRIM8 regulates EWS-FLI1 protein degradation in Ewing sarcoma and its depletion results in EWS/FLI1-mediated oncogene overdose, driving DNA damage and apoptosis of tumor cells [Seong et al., 2021]. Thus, as an E3 ubiquitin ligase itself, TRIM8 presents a unique target for therapeutic degradation. Finally, KRAS is the most frequently mutated oncoprotein, occurring in over 25% of all cancer patients. Due to its smooth and shallow surface, it is considered largely undruggable by standard small molecules, and its structure is evasive due to its conformational disorder as a transcription factor protein [Huang et al., 2021].

We searched existing PPI databases to identify putative interacting partners of our three targets: ACE2 for RBD, PIAS3 for TRIM8, and RAF1 for KRAS [Szklarczyk et al., 2020]. We input these pairs into both our Cut&CLIP pipeline, as well as a co-folding pipeline that adapts the AlphaFold-Multimer complex prediction algorithm followed by PeptiDerive (AF2-CoFold+PeptiDerive) [Evans et al., 2021, Sedan et al., 2016]. After candidate peptide derivation, we experimentally cloned plasmids expressing eight peptides of variable lengths (*≤* 18 amino acids) for each target, directly fused to the CHIPΔTPR uAb domain via a short glycine-serine linker (GSGSG). We subsequently co-transfected these vectors into human HEK293T cells alongside plasmids expressing the target protein fused to superfolder green fluorescent protein (sfGFP) and analyzed the reduction of GFP+ signal (and thus target degradation) via flow cytometry (Figure 4A). Our results demonstrate that select Cut&CLIP-derived peptides induce robust target degradation for all three targets, even on the “undruggable” KRAS oncoprotein. In comparison, the structure-based strategy, while successfully identifying degraders to RBD and TRIM8, failed to produce constructs that induce > 50% degradation of KRAS (Figure 4B).

**Figure 4:**
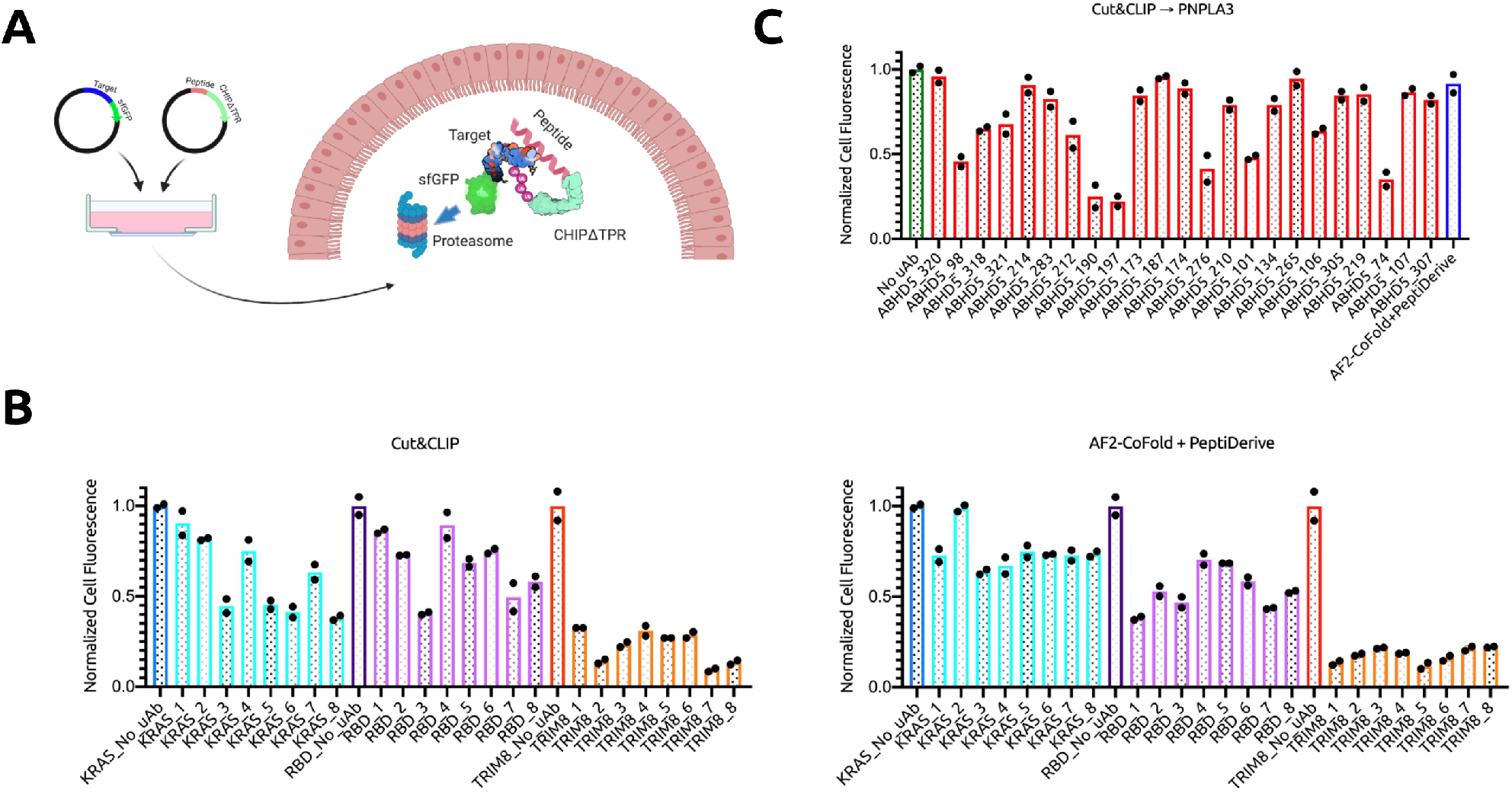
Experimental validation of Cut&CLIP. A) Architecture and mechanism of uAb degradation system. CHIPΔTPR is fused to the C-terminus of targeting peptides, and can thus tag target-sfGFP complexes for ubiquitinmediated degradation in the proteasome, post-plasmid transfection. B) Analysis of KRAS-sfGFP, RBD-sfGFP, and TRIM8-sfGFP degradation via flow cytometry. All samples were performed in independent transfection duplicates (n=2) and gated on GFP+ fluorescence. Normalized cell fluorescence was calculated by dividing the %GFP+ of samples to that of their respective “No uAb” control. C) Analysis of PNPLA3-sfGFP degradation via flow cytometry. All samples were performed in independent transfection duplicates (n=2) and gated on GFP+ fluorescence. Normalized cell fluorescence was calculated by dividing the %GFP+ of samples to that of the “No uAb” control. The final peptide was derived from the AF2-CoFold+PeptiDerive strategy on PNPLA3-ABDH5.

Finally, as uAbs are genetically-encoded constructs, their therapeutic application is limited by the need for *in vivo* delivery vehicles, most of which home to the liver, such as lipid nanoparticles (LNPs) [Hou et al., 2021]. Thus, to extended Cut&CLIP’s utility to a viable therapeutic target, we designed peptides to PNPLA3, a known driver of fatty liver disease, by employing its direct interacting protein, ABHD5 [Yang et al., 2019]. Post transfection and flow cytometry, we show that Cut&CLIP identifies potent ABHD5-derived peptides that enable over 80% degradation of PNPLA3, thus motivating potential clinical translation of our technology (Figure 4C).

## Discussion

Eradicating disease is one of the greatest challenges for the future of human health, and protein-targeting therapeutics have served as potent solutions to this problem. As an example, targeted protein degradation with proteolysis targeting chimeras (PROTACs) and molecular glues utilize small molecules to bind to intracellular proteins transiently and direct their proteolysis by recruiting endogenous E3 ubiquitin ligases [Békés et al., 2022]. More recently, the development of the uAb technology has provided a modular, genetically-encoded alternative to achieve selective degradation of potentially “undruggable” proteins [Portnoff et al., 2014, Chatterjee et al., 2020].

In this work, we exploit recent advancements in contrastive deep learning to design peptides to specified target proteins. Our final models accurately retrieve peptides for known protein-peptide pairs, and more importantly, prioritize candidates that demonstrate effective intracellular target degradation when integrated into the uAb architecture. Our final Cut&CLIP model employs natural binding partners as scaffolds for peptide generation, thus representing a streamlined, efficient, sequence-based pipeline to generate degraders to diverse proteins in the proteome.

In the past few years, protein structure prediction has experienced a wave of excitement with the advent of AlphaFold2 [Jumper et al., 2021]. With these prediction methods in hand, the protein design community now possesses new tools to generate custom proteins with enhanced or novel functionality [Anishchenko et al., 2021, Cao et al., 2022]. In this work, we provide a use-case for which methods like AlphaFold2 may be inferior to pure sequence-based models like Cut&CLIP. Though the AF2-CoFold+PeptiDerive pipeline described in this study managed to produce viable degraders, it struggled to predict large and disordered protein complexes, highlighting its main drawback: efficiency. To generate TRIM8-targeting peptides from PIAS3, the AF2-CoFold+PeptiDerive pipeline required 3 hours, 17 minutes, and 50 seconds on a powerful AWS p3.2xlarge instance with 8 CPU cores and a Nvidia V100 GPU, resources to which many researchers do not have access. Cut&CLIP, on the other hand, only required 15 minutes and 58 seconds for the equivalent design task (including receptor MSA generation) on a standard machine with 2 CPU cores, 8 GB of RAM, and no GPU. Additionally, while both models produced highly effective peptides for TRIM8 and RBD, only Cut&CLIP produced effective degraders for one of the most challenging cancer targets, KRAS.

To further contextualize the power of contrastive sequence-based models for protein design and screening, the model results shown here are based upon the strong assumption that within a batch of 250 peptides, only one is a viable binder. In most applications, especially when using a known interacting partner as a scaffold for peptide generation, there are likely multiple candidates that bind to the queried target. Our experimental results support this observation, as potent degraders were identified by only testing 8 candidates for KRAS, RBD, and TRIM8.

Furthermore, just as OpenAI’s CLIP joint image-text embedding space enables generative models like DALL-E 2 to generate images conditioned on a caption embedding, our joint receptor-peptide embedding space naturally indicates opportunities for *de novo* generation of peptides [Ramesh et al., 2022]. Once we have a target protein embedding, the optimal peptide embedding is known, and all that needs to be done is to decode the peptide embedding back into a sequence. This could be especially critical for generating degraders to targets where minimal experimental evidence exists of potential interacting partners. Considering this versatility, the CLIP architecture thus serves as an ideal framework for target-specified peptide design, as compared to more standard models.

Overall, this work represents an initial application of sequence-based language models to therapeutically-relevant protein design. Future iterations of Cut&CLIP, for example, will incorporate *K*_*d*_ values for high-affinity peptide design and predict the off-targeting propensity of generated sequences. Most importantly, by integrating Cut&CLIP and uAb technology with effective delivery vehicles, such as adeno-associated vectors (AAVs) or LNPs, the peptide-guided protein degradation platform presented here may eventually serve as a potent therapeutic strategy to address a host of diseases deemed untreatable by standard small molecule-based means.

## Methods

### Resources and Model Evaluation

Model architecture, loss functions, and inference methodologies are described in the Results section. Models were trained on a single Nvidia V100 GPU with 32 GB VRAM, as well as 10 CPU cores with a total of 90 GB of RAM. Top-*k* accuracy was calculated to be the probability that the correct peptide is in the top *k* when provided a fixed batch of 250 candidate peptides. Peptide inference was conducted on a machine with 2 CPU cores and 8 GB of RAM.

### Generation of Plasmids

pcDNA3-SARS-CoV-2-S-RBD-sfGFP (Addgene #141184) and pcDNA3-R4-uAb (Addgene #101800) were obtained as gifts from Erik Procko and Matthew DeLisa, respectively. Target coding sequences (CDS) were synthesized as gBlocks from Integrated DNA Technologies (IDT). Sequences was amplified with overhangs for Gibson Assembly-mediated insertion into the pcDNA3-SARS-CoV-2-S-RBD-sfGFP backbone linearized by digestion with NheI and BamHI. An Esp3I restriction site was introduced immediately upstream of the CHIPΔTPR CDS and GSGSG linker via the KLD Enzyme Mix (NEB) following PCR amplification with mutagenic primers (Genewiz). For peptide CDS assembly, oligos were annealed and ligated via T4 DNA Ligase into the Esp3I-digested uAb backbone. Assembled constructs were transformed into 50 *μ*L NEB Turbo Competent *Escherichia coli* cells, and plated onto LB agar supplemented with the appropriate antibiotic for subsequent sequence verification of colonies and plasmid purification.

### Cell Culture and Flow Cytometry

HEK293T cells were maintained in Dulbecco’s Modified Eagle’s Medium (DMEM) supplemented with 100 units/ml penicillin, 100 mg/ml streptomycin, and 10% fetal bovine serum (FBS). Target-sfGFP (50 ng) and peptide-CHIPΔTPR (50 ng) plasmids were transfected into cells as duplicates (2×10^4^/well in a 96-well plate) with Lipofectamine 3000 (Invitrogen) in Opti-MEM (Gibco). After 3 days post transfection, cells were harvested and analyzed on a FACSCelesta for GFP fluorescence (488-nm laser excitation, 530/30 filter for detection). Cells expressing GFP were gated as compared to a GFP-control, and normalized cell fluorescence was calculated to the “No uAb” control.

### Statistics and Reproducibility

All samples were performed in independent transfection duplicates (n=2), and normalized cell fluorescence values were averaged.

## Declarations

## Acknowledgements

We thank the MIT SuperCloud and Duke Compute Cluster for providing the HPC and database resources that have contributed to the research reported within this manuscript. We further thank Tianzheng Ye and Dr. Matthew Delisa for uAb plasmids, Dr. Joseph Jacobson for conceptual input, and Dr. Neil Gershenfeld and Dr. Shuguang Zhang for shared lab equipment.

## Author Contributions

P.C. conceived, designed, directed, and supervised the study. K.P. and M.P. curated peptide-protein data. K.P. conceived, designed, and implemented the Cut&CLIP workflow, with assistance from S.B. and G.B. M.P. and S.B. conceived, designed, and implemented the AF2-CoFold+PeptiDerive workflow. E.T., T.S., and S.R.T.K. built constructs, conducted transfections, carried out degradation experiments, and performed data analyses. P.C. and K.P. wrote the paper, with input from all authors.

## Data and Materials Availability

All data needed to evaluate the conclusions in the paper are present in the paper and tables. All peptide sequences, source data, and results can be found at: https://tinyurl.com/cutnclip. Cut&CLIP code will be made available on GitHub upon publication. uAb expression plasmids will be deposited to Addgene.

